# Direct and indirect interactions promote complexes of the lipoprotein LbcA, the CtpA protease and its substrates, and other cell wall proteins in *Pseudomonas aeruginosa*

**DOI:** 10.1101/2021.07.29.454410

**Authors:** Dolonchapa Chakraborty, Andrew J. Darwin

## Abstract

The *Pseudomonas aeruginosa* lipoprotein LbcA was discovered because it copurified with and promoted the activity of CtpA, a carboxyl-terminal processing protease (CTP) required for type III secretion system function, and for virulence in a mouse model of acute pneumonia. In this study we explored the role of LbcA by determining its effect on the proteome and its participation in protein complexes. *lbcA* and *ctpA* null mutations had strikingly similar effects on the proteome, suggesting that facilitating CtpA might be the most impactful role of LbcA in the bacterial cell. Independent complexes containing LbcA and CtpA, or LbcA and substrate, were isolated from *P. aeruginosa* cells, indicating that LbcA facilitates proteolysis by recruiting the protease and its substrates independently. An unbiased examination of proteins that copurified with LbcA revealed an enrichment for proteins associated with the cell wall. One of these copurification partners was found to be a new CtpA substrate, and the first substrate that is not a peptidoglycan hydrolase. Many of the other LbcA copurification partners are known or predicted peptidoglycan hydrolases. However, some of these LbcA copurification partners were not cleaved by CtpA, and an *in vitro* assay revealed that while CtpA and all of its substrates bound to LbcA directly, these non-substrates did not. Subsequent experiments suggested that the non substrates might co-purify with LbcA by participating in multi-enzyme complexes containing LbcA-binding CtpA substrates.

**IMPORTANCE:** Carboxyl-terminal processing proteases (CTPs) are widely conserved and associated with the virulence of several bacteria, including CtpA in *Pseudomonas aeruginosa*. CtpA copurifies with the uncharacterized lipoprotein, LbcA. This study shows that the most impactful role of LbcA might be to promote CtpA-dependent proteolysis, and that it achieves this as a scaffold for CtpA and its substrates. It also reveals that LbcA copurification partners are enriched for cell wall-associated proteins, one of which is a novel CtpA substrate. Some of the other LbcA copurification partners are not cleaved by CtpA, but might copurify with LbcA because they participate in multi-enzyme complexes containing CtpA substrates. These findings are important, given the links between CTPs, their associated proteins, peptidoglycan remodeling, and virulence.

## INTRODUCTION

*Pseudomonas aeruginosa* is a Gram-negative bacterium that is widespread in the environment and responsible for serious opportunistic infections, especially in healthcare settings (1). As with any bacterial pathogen, the cell envelope of *P. aeruginosa* is the focal point for its interaction with a host. Many critical virulence factors are assembled or exported through the cell envelope, including the type III secretion system (T3SS) and the polysaccharide alginate, which play important roles in acute and chronic infections, respectively (2, 3). The peptidoglycan cell wall is a layer of the cell envelope that helps maintain bacterial cell integrity, has to be remodeled during growth and virulence factor assembly/export, and is one of the most important targets for antibiotics (4–6). Therefore, a comprehensive understanding of proteins involved in cell wall assembly, remodeling and associated functions has obvious significance.

The cleavage of proteins and peptides in the cell envelope plays roles in protein export, protein quality control and turnover, signal transduction, and the integrity and functions of the cell wall (7–10). One widely conserved family of serine proteases that function within the bacterial cell envelope is the carboxyl-terminal processing proteases (CTPs). Their name arose from early findings that CTPs can process protein substrates to a functional form by removing a small segment from their C-terminal end (11–13). However, more recent findings have demonstrated that at least one CTP removes an N-terminal segment from its substrate (14). It has also emerged that bacterial CTPs can completely degrade their substrates, including some peptidoglycan cross-link hydrolases, and at least one lytic transglycosylase, which are degraded by CTPs in *Escherichia coli* and *P. aeruginosa* (15–18). Therefore, CTPs appear to be one of the mechanisms used by bacteria to control cell wall hydrolase activity.

Another feature that has been uncovered in the last few years, for two quite different CTPs, is that they can function in partnership with another protein. The *E. coli* Prc protease interacts with the tetratricopeptide repeat (TPR)-containing outer membrane lipoprotein NlpI to degrade the NlpC/P60 endopeptidase MepS, which is a peptidoglycan cross-link hydrolase (16). Structural analysis suggests that NlpI recruits MepS and helps to feed it into the Prc active site for destruction (19). However, Prc can also cleave at least three substrates without the help of NlpI (15, 18, 20). In *P. aeruginosa*, the CtpA protease interacts with the TPR-containing lipoprotein LbcA to degrade four different predicted peptidoglycan cross-link hydrolases, the NlpC/P60 endopeptidases PA1198 and PA1199, and the LytM/M23 endopeptidases MepM and PA4404 (17). Despite the obvious similarities between the NlpI-Prc and LbcA-CtpA systems, it is not at all clear if they function by the same molecular mechanisms. *E. coli* Prc and *P. aeruginosa* CtpA are not orthologs, being quite different in size and belonging to different CTP subfamilies (21–23). In fact, *P. aeruginosa* has a second CTP that is the apparent ortholog of *E. coli* Prc (24–26). In addition to the differences between the *E. coli* Prc and *P. aeruginosa* CtpA proteases, although their NlpI and LbcA lipoprotein partners both have the short degenerate TPR repeat motifs, they are very different in size and share no obvious primary sequence homology when aligned (17).

In this study we have investigated the role of LbcA in *P. aeruginosa*. We found that *lbcA* and *ctpA* null mutations have remarkably similar effects on the proteome, suggesting that the most impactful role of LbcA might be to facilitate CtpA-dependent proteolysis. LbcA does this, at least in part, by binding to CtpA and its substrates directly, but independently, to bring them together for proteolysis. We also found that the most abundant LbcA copurification partners are enriched for several cell-wall associated proteins, and that one of these proteins is a newly discovered CtpA substrate. Some other LbcA copurification partners are not CtpA substrates, and unlike the substrates, our data suggests that they might associate with LbcA indirectly by participating in cell wall enzyme complexes.

## RESULTS

### *lbcA* and *ctpA* null mutations have similar effects on the proteome

To begin to investigate the role of LbcA, we used label-free quantitative proteomics (LFQP) to compare protein levels in wild type, Δ*ctpA*, and Δ*lbcA* strains. When *p* values were determined and a 5% Benjamini-Hochberg based false discovery rate cutoff was applied, many proteins had significantly different levels when either the Δ*ctpA* or Δ*lbcA* mutant was compared to the wild type, but not when the two mutants were compared to each other (Fig. 1, Supplemental Tables 1-3). This suggests that Δ*ctpA* and Δ*lbcA* mutations had broadly overlapping effects on the proteome. One caveat is that the amount of variation between the triplicate samples of each individual strain affects the ability to make statistically significant calls between strains. However, a closer examination of the data also supported the conclusion. Two CtpA substrates, MepM and PA1198, could be quantified in all three strains, and were within the top 25 proteins ranked by fold-increase in abundance when either the Δ*ctpA* or Δ*lbcA* mutant was compared to wild type (Fig. 1, Supplemental Tables 1-2). Other changes in the mutants were also similar, and many might be caused by effects on gene expression. For example, PA0634 had the largest increase in abundance in the Δ*ctpA* and Δ*lbcA* mutants, and several other proteins encoded by genes closely linked to PA0634 also increased (Fig.1; Supplemental Tables 1-2). This region encodes the R- and F-type pyocins, which are induced by stresses including DNA-damage and overproduction of extracytoplasmic function sigma factor PA4896 (27, 28). Stress from the accumulating substrates of CtpA might induce pyocin gene expression indirectly. There was also overlap in the most reduced abundance proteins in the Δ*ctpA* and Δ*lbcA* mutants, which included some regulators, components and effectors of the type III secretion system (T3SS; Supplemental Tables 1-2). Although our experiments were not done in T3SS-inducing conditions, this is consistent with Δ*ctpA* compromising T3SS function (23). The only striking difference between the Δ*ctpA* and Δ*lbcA* mutants was increased abundance of LolB and Ipk in the Δ*lbcA* mutant only (Fig. 1). However, this is a trivial consequence of *lbcA* being replaced by the gentamycin resistance gene *aacC1* in the same orientation as *lbcA*, in order to maintain sufficient expression of the essential downstream genes *lolB* and *ipk* (17). Therefore, these LFQP data support the hypothesis that a major role of LbcA is to enable CtpA-dependent proteolysis.

**FIG 1.**
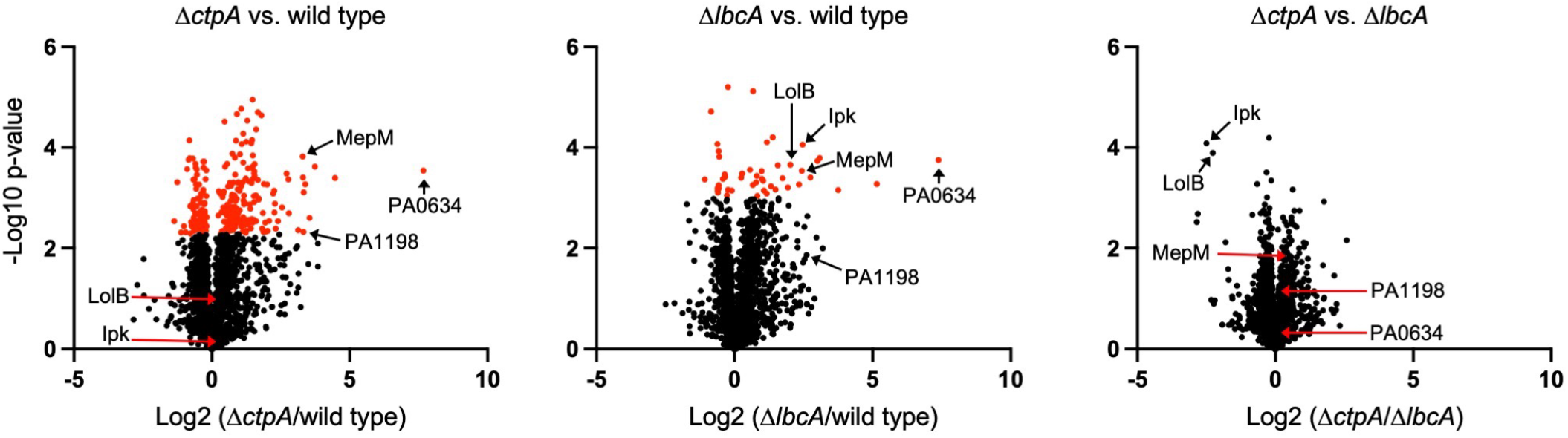
Δ*lbcA* and Δ*ctpA* mutants have similar effects on the proteome. Each chart shows the relative protein levels (x-axis) between the pair of strains indicated above, and the statistical significance of each difference (y-axis). Circular symbols represent individual proteins. Data for the CtpA and LbcA proteins themselves are not included in the plots. Symbols in red indicate proteins for which their different level in the comparison was considered significant after a 5% Benjamini-Hochberg based false discovery rate cutoff was applied.

### LbcA can form independent complexes with CtpA and its substrates

LbcA has several tetratricopeptide (TPR) motifs, which mediate protein-protein interactions and the formation of multi-protein complexes (17, 29). Therefore, we hypothesized that LbcA enables CtpA-dependent proteolysis by acting as a scaffold protein to recruit CtpA and its substrates independently. To test this, we used co-immunoprecipitation (co-IP) assays to investigate *in vivo* complexes containing LbcA, CtpA and the representative CtpA substrate, PA1198.

The first bait was LbcA with a C-terminal FLAG tag (LbcA-FLAG), which was produced in wild type, Δ*ctpA*, *ctpA*-*S302A* or ΔPA1198 strains. The *ctpA*-*S302A* mutation destroys the proteolytic activity of CtpA, allowing analysis of complexes that contain an intact substrate (17). Analysis of anti-FLAG monoclonal antibody immunoprecipitates revealed that both CtpA and PA1198 copurified with LbcA-FLAG in the *ctpA-S302A* strain (Fig 2A). PA1198 still copurified with LbcA-FLAG in a Δ*ctpA* mutant, and CtpA still copurified with LbcA-FLAG in a ΔPA1198 mutant. Therefore, these data suggest that LbcA can form independent complexes with CtpA and its substrate PA1198 *in vivo*. As a specificity control, the unrelated periplasmic PA0943 protein (30) was not detected by immunoblot analysis of LbcA-FLAG immunoprecipitates from any strain. Furthermore, when the LbcA bait did not have a FLAG tag, none of the proteins tested were detected above trace levels in anti-FLAG immunoprecipitates (Fig 2A).

**FIG 2.**
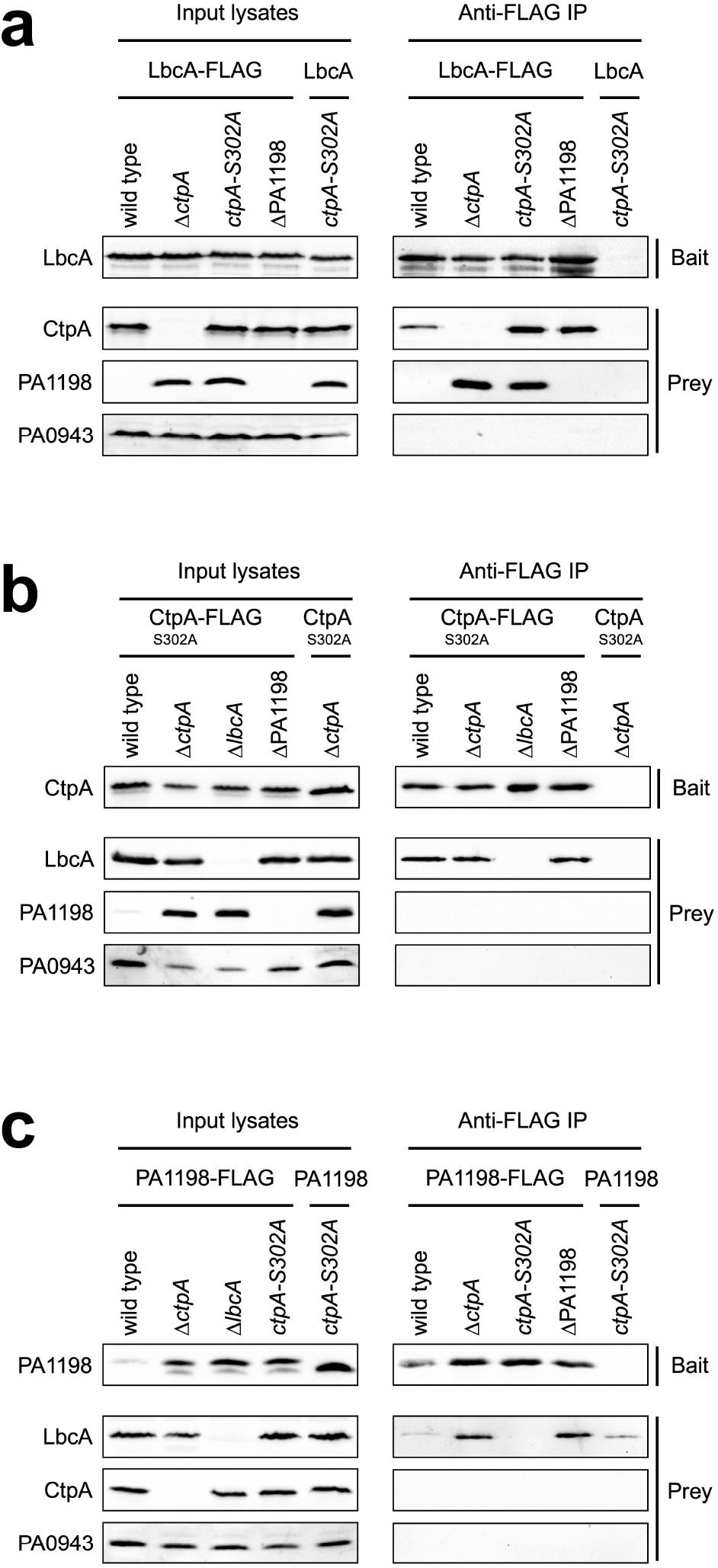
LbcA can form independent complexes with CtpA and it substrates. Each panel shows immunoblot analysis of input lysates and immunoprecipitates (Anti-FLAG IP). Strain genotypes are shown above each individual lane. Polyclonal antisera used for detection are shown at the left. a) Baits were LbcA-FLAG or LbcA negative control. b) Baits were CtpA-S302A-FLAG-His_6_ or CtpA-S302A-His_6_ negative control. c) Baits were PA1198-FLAG or PA1198 negative control.

Next, we used CtpA-S302A with a C-terminal FLAG-His_6_ tag as the bait protein (17). Analysis of the anti-FLAG immunoprecipitates revealed that LbcA copurified with CtpA, but the PA1198 substrate did not (Fig. 2B). This further supports the presence of an LbcA•CtpA complex, but it suggests that CtpA cannot form a stable complex with its substrate alone. Indeed, when we went on to use the PA1198-FLAG substrate as the bait, LbcA copurified with PA1198, but CtpA did not (Fig. 2C). Therefore, all of these co-IPs suggest that LbcA can form independent complexes with CtpA or its substrate, but CtpA cannot form a stable binary complex with its substrate alone. This supports the hypothesis that LbcA functions, at least in part, by binding to CtpA and its substrate independently, in order to bring them together for proteolysis.

### LbcA copurification partners are enriched for cell wall enzymes and binding proteins

The preceding experiments suggested that a major role of LbcA is to facilitate CtpA-dependent proteolysis, which it achieves by acting as a scaffolding protein. Therefore, we reasoned that another unbiased approach to learn more about the targets of the LbcA•CtpA complex, and perhaps any additional role(s) of LbcA, would be to identify LbcA binding partners. For this, two independent purifications of LbcA-FLAG from a *ctpA-S302A* strain were analyzed by mass spectrometry (Supplemental Table 4). To control for specificity, we also analyzed two independent purifications in which the LbcA bait protein was not FLAG-tagged (Supplemental Table 5). 67 known or predicted envelope proteins were enriched in the LbcA-FLAG purification (Supplemental Table 6). CtpA was the most abundant of these proteins based on peptide spectral matches (PSMs), and three of the known CtpA substrates, MepM, PA1198 and PA1199, were also detected. A striking observation was that half of the 18 most abundant proteins that copurified with LbcA-FLAG were known or predicted cell wall associated proteins (Supplemental Table 6). This is in contrast to only 5 of the remaining 49 lower abundance proteins. Therefore, the most abundant LbcA copurification partners were enriched for cell wall enzymes and cell wall binding proteins. These data suggest that LbcA is associated with multiple cell wall-related proteins, beyond the previously known proteolytic substrates of CtpA.

### Proteins in complex with LbcA *in vivo* include CtpA substrates and non-substrates

We could not follow up on all 67 envelope proteins that were enriched in the LbcA-FLAG immunoprecipitation (Supplemental Table 6). However, one of our motivations was to identify new substrates of the LbcA•CtpA complex. In a previous study, the CtpA substrates MepM and PA1198 were discovered because they were the most abundant proteins trapped by an inactive LbcA•CtpA-S302A complex after *in vivo* cross-linking (17). Many other proteins were trapped exclusively by the inactive LbcA•CtpA-S302A complex, but again they were too numerous for individual follow up. Therefore, for this study, out of the 67 proteins that co-purified with LbcA-FLAG, we decided to focus on those that had also been trapped by the inactive LbcA•CtpA-S302A complex exclusively in the previous work (17). The addition of this criterion reduced the number of LbcA-FLAG co-purification partners for follow up to seven proteins, all of which are predicted cell wall hydrolases or peptidoglycan-binding proteins (Table 1). These included the two known CtpA substrates MepM and PA1198, but also three proteins, PA1048, MltB1 and AmiB, which had not been investigated before for the possibility of a relationship with LbcA and/or CtpA. The remaining two proteins, the lytic transglycosylases RlpA and MltD, were assessed as possible CtpA substrates in the previous study. However, RlpA-FLAG and MltD-FLAG did not accumulate in a Δ*ctpA* mutant, suggesting that they are not CtpA substrates (17). Nevertheless, we included RlpA and MltD in our follow up here because the previous analysis was not exhaustive.

**TABLE 1.** Envelope proteins that copurified with LbcA and were also isolated by a LbcA•CtpA-S302A protease trap ^*a*^

| Protein | Predicted features/function | Abundance rank<br>with LbcA <sup>b</sup> | Abundance rank<br>with LbcA•CtpA trap <sup>c</sup> | Srivastava *et al.*<br>CtpA substrate <sup>d</sup> |
| --- | --- | --- | --- | --- |
| PA1198 | Peptidoglycan NlpC/P60 endopeptidase | 3 | 2 | Yes |
| PA1048 | Peptidoglycan-binding OmpA C-terminal domain | 4 | 7 | NT |
| RlpA | Peptidoglycan lytic transglycosylase | 5 | 4 | No |
| MltD | Peptidoglycan lytic transglycosylase | 10 | 3 | No |
| MltB1 | Peptidoglycan lytic transglycosylase | 13 | 6 | NT |
| MepM | Peptidoglycan LytM/M23 endopeptidase | 18 | 1 | Yes |
| AmiB | Peptidoglycan amidase | 65 | 23 | NT |

<sup>a</sup> Cell envelope proteins that co-purified with LbcA•CtpA-S302A, but not with LbcA•CtpA, following *in vivo* cross-linking (17), and also co-purified with LbcA-FLAG from a *ctpA-S302A* strain in this study with average peptide spectral matches $\geq 5$ - fold higher than in a LbcA no FLAG tag negative control purification (Supplemental Tables 4-6).
<sup>b</sup> Rank based on average number of peptide spectral matches amongst 67 cell envelope proteins that co-purified with LbcA-FLAG (Supplemental Table 6)
<sup>c</sup> Rank based on average number of peptide spectral matches amongst 25 cell envelope proteins that co-purified exclusively with the LbcA•CtpA-S302A trap (17).
<sup>d</sup> In Srivastava *et al.*, a protein was classified as a CtpA substrate if a FLAG-tagged version accumulated in a $\Delta ctpA$ mutant *in vivo* and the protein was degraded by CtpA *in vitro* (17). NT = not tested in Srivastava *et al.*

We tested rigorously if these proteins might be CtpA substrates, with the previously characterized substrates MepM and PA1198 serving as positive controls. To screen *in vivo*, we constructed plasmids encoding C-terminal FLAG tagged versions, expressed from a non-native promoter, and determined if they accumulated in a Δ*ctpA* mutant (Fig. 3a). As reported previously, the CtpA substrates MepM and PA1198 accumulated in the Δ*ctpA* mutant, whereas RlpA and MltD did not (17). MltB1 and AmiB did not accumulate in the Δ*ctpA* mutant, suggesting that like RlpA and MltD, they are not CtpA substrates. However, PA1048 behaved like the known CtpA substrates. It was barely detectable in the wild type, but accumulated robustly in the Δ*ctpA* mutant (Fig. 3a). We also examined the data for these proteins from our earlier LFQP analysis and found that the endogenous proteins had a similar pattern to the plasmid encoded FLAG-tagged versions. MepM, PA1198 and PA1048 levels were approximately 6- to 10-fold higher in the Δ*ctpA* and Δ*lbcA* mutants, whereas MltD, RlpA and AmiB had more modest changes (Fig. 3b; note that in the LFQP analysis MltB1 was not detected well in multiple samples and so could not be quantified).

**FIG 3.**
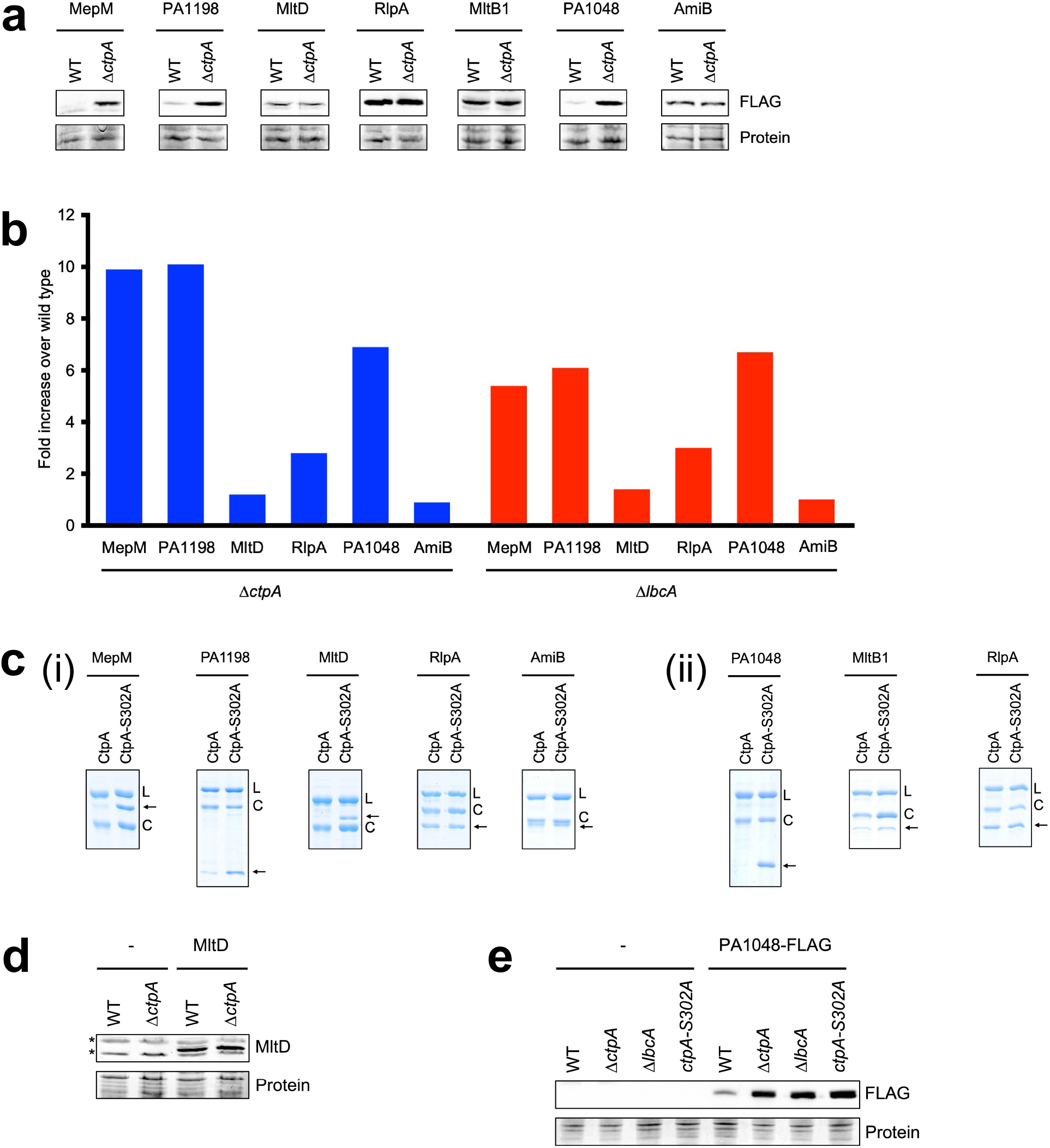
Analysis of LbcA copurification partners. a) Immunoblots of whole-cell lysates of wild-type (WT) and Δ*ctpA* strains containing a plasmid encoding C-terminal FLAG-tagged versions of the indicated proteins. Proteins were detected with anti-FLAG monoclonal antibodies (FLAG). b) LFQP analysis of endogenous proteins in Δ*ctpA* and Δ*lbcA* mutants. c) *In vitro* proteolysis assay. Natively purified (i) or renatured (ii) N-terminal His_6_-tagged versions of the proteins indicated at the top were incubated with LbcA-His_6_ and either CtpA-His_6_ or CtpA-S302A-His_6_ for 3 h at 37°C. Samples were separated on SDS-PAGE gels and stained with Coomassie brilliant blue. L = LbcA-His_6_; C = CtpA-His_6_ or CtpA-S302A-His_6_; arrows indicate the N-terminal His_6_-tagged protein being tested. d) Immunoblot analysis of whole-cell lysates of wild-type (WT) and Δ*ctpA* strains containing a plasmid encoding untagged MltD or the empty vector (−). MltD was detected with a polyclonal antiserum. All strains had a Δ*mltD* mutation. Asterisks indicate cross-reactive proteins. e) Immunoblot analysis of whole-cell lysates of wild-type (WT), Δ*ctpA*, Δ*lbcA* or *ctpA-S302A* strains containing a plasmid encoding PA1048-FLAG or the empty vector (−). For panels a, d and e, loading was determined by a Ponceau S total protein stain of the nitrocellulose membrane used for detection (Protein).

**FIG 4.**
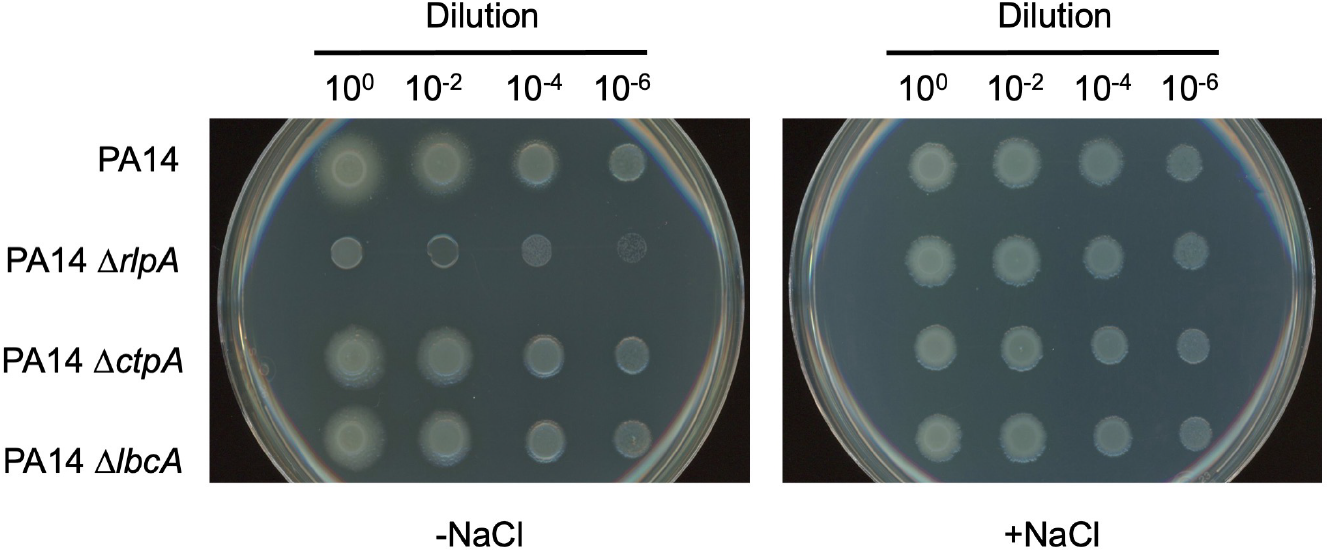
Evidence that LbcA or CtpA are not essential for RlpA function. Serial dilutions of normalized saturated cultures were spotted onto LB agar without NaCl (−NaCl) or containing 1% (w/v) NaCl (+ NaCl) and incubated at 37°C for approximately 16 h.

Next, we tested if N-terminal His_6_-tagged versions of these proteins could be cleaved by CtpA *in vitro*, as described previously (17). In agreement with the *in vivo* data, MepM and PA1198 were degraded, whereas RlpA and AmiB were not, which confirmed that RlpA and AmiB are not CtpA substrates (Fig. 3c). However, MltD was degraded *in vitro*, even though MltD-FLAG and endogenous MltD did not accumulate in Δ*ctpA* mutants *in vivo*. This left open the possibility that MltD is a CtpA substrate (see below). PA1048 and MltB1 were insoluble when overproduced in *E. coli* prior to purification. To circumvent this, we solubilized PA1048 and MltB1 by denaturation and then renatured them on a nickel agarose column prior to elution. To ensure that this would not render any protein a CtpA substrate, we also repurified RlpA in the same way. Analysis of these renatured proteins showed that PA1048 was degraded by CtpA, whereas MltB and the RlpA control were not (Fig. 3c). Therefore, together all of these analyses showed that the LbcA copurification partners RlpA, AmiB, MltB1, and possibly MltD, are not CtpA substrates.

### PA1048 is a novel substrate of CtpA

We investigated PA1048 and MltD further because the previous analyses suggested that PA1048 is a new CtpA substrate, but were inconclusive for MltD, which was degraded *in vitro* but did not accumulate *in vivo* in a Δ*ctpA* mutant (Fig. 3a-c). It was possible that endogenous MltD production was downregulated in a Δ*ctpA* mutant, which could obscure detection of decreased degradation in LFQP analysis. This was not a concern for the plasmid encoded MltD-FLAG, but in this case, the C-terminal FLAG tag could have interfered with MltD degradation. To address these issues, we raised a polyclonal MltD antiserum and used it to detect untagged MltD encoded on a plasmid with a non-native promoter. This confirmed that MltD does not accumulate in a Δ*ctpA* mutant (Fig. 3d). Therefore, MltD is not a CtpA substrate *in vivo*, at least under the growth conditions used in this study (see Discussion).

Our analysis had suggested that PA1048 is a newly discovered CtpA substrate because PA1048-FLAG accumulated in a Δ*ctpA* strain, the level of endogenous PA1048 was much higher in Δ*ctpA* and Δ*lbcA* strains, and His_6_-PA1048 was degraded by LbcA•CtpA *in vitro* (Fig. 3abc). However, to ensure that PA1048 behaves exactly like all previously known substrates, we also tested the effects of Δ*lbcA* and *ctpA-S302A* mutations on constitutively produced PA1048 *in vivo*. PA1048-FLAG accumulated indistinguishably in Δ*ctpA*, Δ*lbcA* and *ctpA-S302A* strains (Fig. 3e). Therefore, PA1048 is the fifth substrate of CtpA to be discovered, because it accumulates in all CtpA protease-defective strains *in vivo* and is degraded by the LbcA•CtpA complex *in vitro*. Unlike the other four CtpA substrates, MepM, PA1198, PA1199 and PA4404, PA1048 is not a predicted peptidoglycan cross-link hydrolase. It is a predicted outer membrane (OM) lipoprotein, with an N-terminal domain of unknown function, and an OmpA-like C-terminal domain that binds to peptidoglycan non-covalently (see Discussion).

### LbcA is not essential for RlpA function

One of the proteins that copurified with LbcA, but is not a CtpA substrate, is RlpA (Figs. 2–3). We considered the possibility that a complex containing LbcA and RlpA might occur because LbcA plays a role in facilitating RlpA function. A *P. aeruginosa rlpA* null mutant was reported to have a viability defect when plated on LB agar lacking NaCl (31). Therefore, we reasoned that we could use that phenotype to test if LbcA is required for RlpA function. However, when we introduced an *rlpA* in frame deletion mutation into the PAK strain used in our studies, the mutant grew normally on LB agar lacking NaCl (data not shown). We suspected that this might be due to the strain background, because the original Δ*rlpA* phenotype was reported in strain PA14 only (31). Therefore, we introduced *rlpA*, *lbcA* and *ctpA* deletion mutations into strain PA14. In this strain background, the Δ*rlpA* mutant had the previously reported viability defect when plated onto LB agar without NaCl (Fig. 5). However, Δ*lbcA* or Δ*ctpA* mutants did not have the same phenotype, which suggests that any interaction between LbcA and RlpA is not essential for RlpA function (Fig. 5). We also considered the reverse possibility, that RlpA might play a role in the functioning of the LbcA•CtpA proteolytic complex. However, a Δ*rlpA* mutation had no effect on the levels of CtpA substrates *in vivo*, suggesting that RlpA does not promote or inhibit LbcA•CtpA proteolytic function (data not shown).

**FIG 5.**
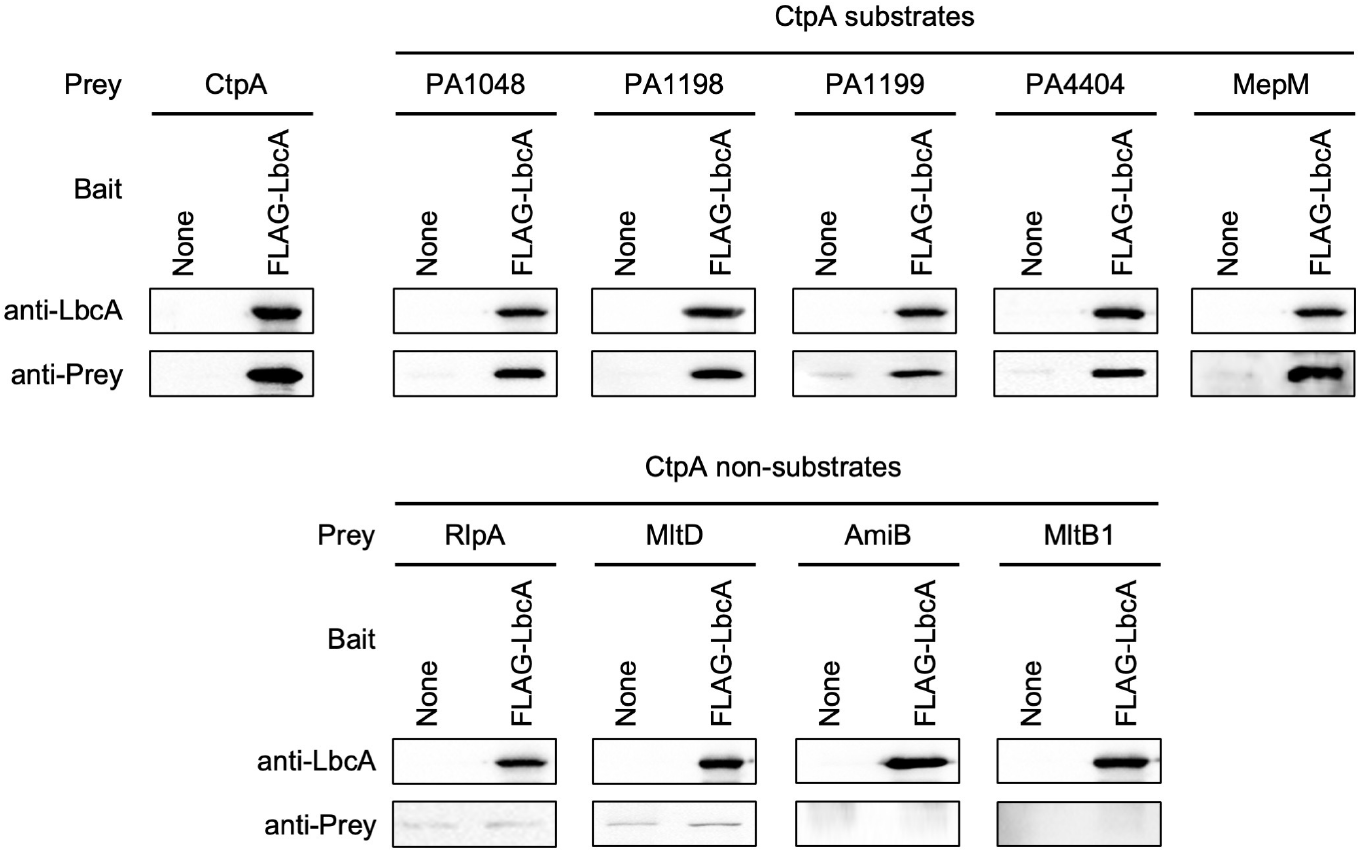
Only CtpA and its substrates bind to LbcA directly *in vitro*. Anti-FLAG M2 affinity resin was used to purify protein from *E. coli* lysates containing the empty pET-24b(+) vector (None) or a derivative encoding FLAG-LbcA. The purified anti-FLAG M2 agarose immunocomplexes were then incubated with approximately 1.5 μg of a purified prey protein (Prey) for 1 h, washed extensively in high stringency buffer containing 0.5 M NaCl and 0.1% (w/v) Triton-X-100, and analyzed by immunoblot.

### Only CtpA and its substrates bind to LbcA directly

Our experiments had revealed that LbcA can form a complex with CtpA and its substrates independently *in vivo*, but that some proteins that are not CtpA substrates also copurify with LbcA. However, the co-IP experiments that led to those conclusions cannot distinguish between direct and indirect interactions. If LbcA acts as a scaffold to facilitate CtpA-dependent proteolysis, we hypothesized that CtpA and its substrates should bind to LbcA directly and independently. Some non-CtpA substrates might also bind to LbcA directly, perhaps as part of an undiscovered LbcA function. Therefore, we used an *in vitro* LbcA-binding assay to uncover which proteins can or cannot bind to LbcA directly.

A purified FLAG-LbcA protein was tested for its ability to interact with individually purified prey proteins. Binding reactions and washes were done in buffer containing 0.5 M NaCl and 0.1% (w/v) Triton-X-100, to ensure high stringency. These experiments confirmed that LbcA and CtpA interact directly, and that all five known CtpA substrates bound to LbcA directly as well, including the new PA1048 substrate discovered in this study (Fig. 5). In striking contrast, all four non-CtpA substrates, which we had identified as LbcA-complex members *in vivo* and focused on in this study, were unable to bind to LbcA directly (Fig. 5). Lowering stringency by reducing salt concentration, or using LbcA bait protein with the FLAG tag on the C-terminus rather than the N-terminus, did not alter any of these conclusions (data not shown). These results support the idea that a major role of LbcA is to act as a scaffolding protein that binds to CtpA and its substrates directly, bringing them together for proteolysis. However, they suggest that some non-CtpA substrates that were cross-linked to an inactive LbcA•CtpA-S302A complex in a previous study, and that copurified natively with LbcA in this study, do not bind to LbcA directly. Therefore, the *in vivo* association of these non-CtpA substrates with LbcA might be indirect, perhaps due to their participation in multi-protein complexes that include one or more proteins that can bind to LbcA. We investigated this possibility in our final series of experiments.

### LbcA-independent complexes between CtpA substrates and non-substrates *in vivo*

We used co-IP assays to probe the relationship between LbcA and its copurification partners. We chose PA1198 as a representative CtpA-substrate, and RlpA as a representative non-CtpA substrate. When RlpA-FLAG was the bait, LbcA and PA1198 copurified with it, as expected from the results of our previous experiments (Fig. 6a; detectible amounts of PA1198 copurified only in the strains where it could not be degraded by CtpA). However, PA1198 still copurified with RlpA-FLAG in a Δ*lbcA* mutant. Therefore, scaffolding by LbcA is not the explanation for RlpA and PA1198 copurification, and this is consistent with the inability of LbcA and RlpA to interact directly (Fig. 5). It also raised the possibility that interactions between CtpA substrates and non-substrates, such as PA1198 and RlpA, might bridge interaction between non-substrates and LbcA. This could explain why proteins such as RlpA and MltD were enriched in the LbcA•CtpA-S302A trap in our previous study (17). However, LbcA still copurified with RlpA-FLAG in a ΔPA1198 strain, showing that bridging by PA1198 cannot be the only explanation (Fig. 6a). These conclusions were corroborated when LbcA-FLAG was the bait. RlpA copurified with LbcA-FLAG in a ΔPA1198 strain, and PA1198 copurified with LbcA-FLAG in a Δ*rlpA* strain (Fig. 6b; all strains had a Δ*ctpA* mutation to ensure similar levels of the endogenous PA1198 prey in all comparisons). The conclusions were also supported with a PA1198-FLAG bait, which copurified with RlpA and LbcA, even in strains that contained only one of them (Fig. 6c).

**FIG 6.**
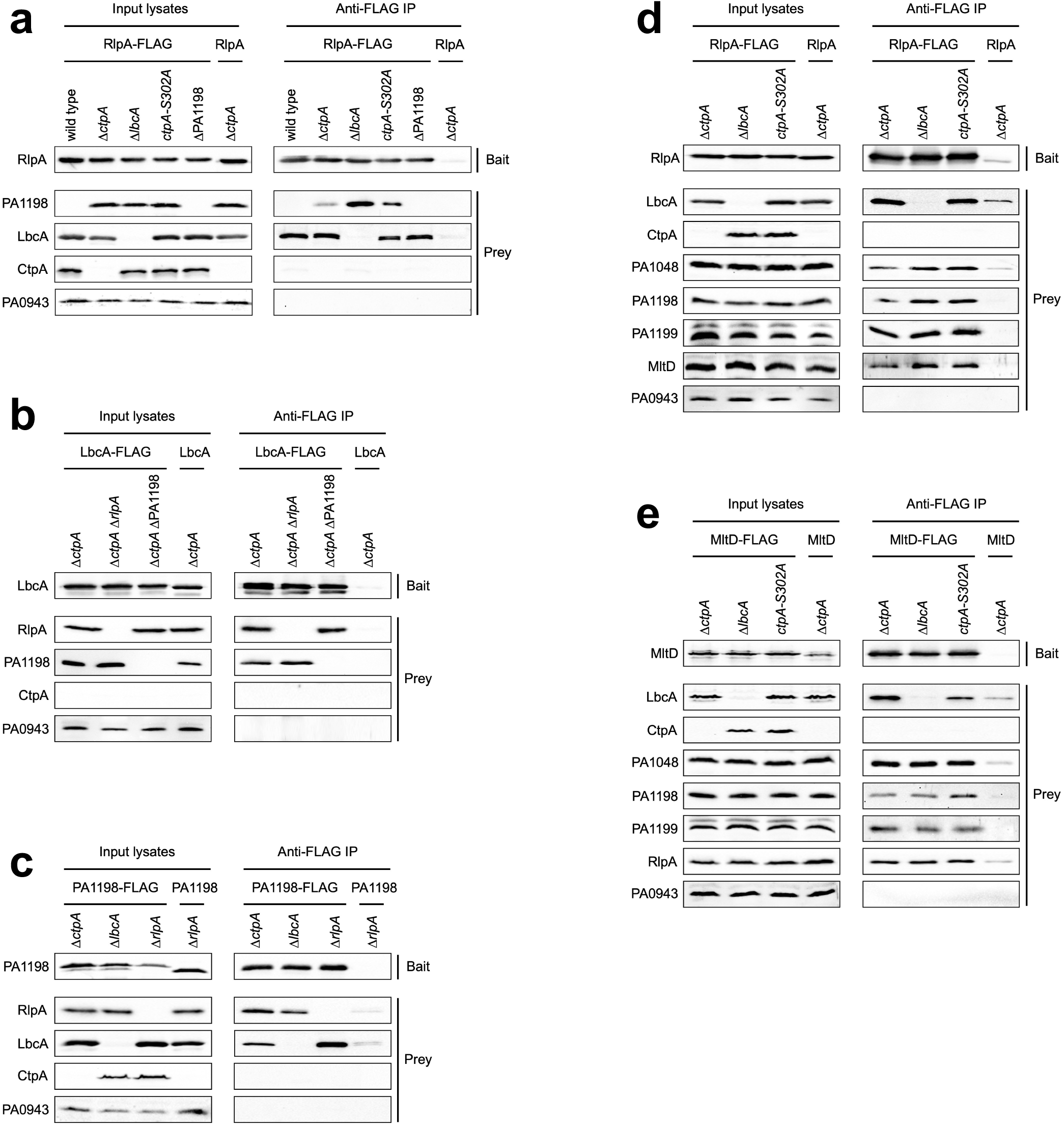
Co-immunoprecipitation analysis suggests the formation of LbcA-independent complexes between CtpA substrates and non-substrates. Each panel shows immunoblot analysis of input lysates and immunoprecipitates (Anti-FLAG IP). Strain genotypes are shown above each individual lane. Polyclonal antisera used for detection are shown at the left. a) Baits were RlpA-FLAG or RlpA negative control. b) Baits were LbcA-FLAG or LbcA negative control. c) Baits were PA1198-FLAG or PA1198 negative control. d) Baits were RlpA-FLAG or RlpA negative control. e) Baits were MltD-FLAG or MltD negative control.

The preceding experiments suggested that LbcA-independent associations between CtpA substrates and non-substrates occur *in vivo*. This could explain, at least in part, why some non-substrates copurified with LbcA and were enriched in an inactive LbcA•CtpA-S302A complex. However, bridging by PA1198 cannot be the sole explanation. Therefore, we expanded the experiments to test if other CtpA substrates and non-substrates copurified with RlpA-FLAG. In addition to PA1198, the CtpA substrates PA1048 and PA1199, and the non-substrate MltD, also co-purified with RlpA-FLAG (Fig. 6d; our polyclonal antisera for the CtpA substrates MepM and PA4404 have low avidity and so we could not reach a conclusion for them). All of these complexes occurred in both *lbcA*^+^ and Δ*lbcA* strains, demonstrating that scaffolding by LbcA was not the explanation. Finally, a similar set of experiments using MltD-FLAG as the bait showed that it copurified with CtpA substrates PA1048, PA1198 and PA1199, and the non-substrate RlpA (Fig. 6e). Once again, all of these complexes were independent of LbcA. Together, all of these experiments suggest that complexes between peptidoglycan hydrolases and other peptidoglycan binding proteins occur in *P. aeruginosa*, and this might explain how some non-CtpA substrates are found in LbcA-containing complexes.

## DISCUSSION

LbcA was discovered because it copurified with CtpA, and it was then found to be required for CtpA-dependent proteolysis of four substrates (17). In this study we combined unbiased approaches with focused experiments to explore the role of LbcA in *P. aeruginosa,* and to gain insight into how it promotes CtpA activity. Our findings suggest that facilitating CtpA-dependent proteolysis might be the most impactful role of LbcA, which it achieves, at least in part, by acting as a scaffold for CtpA and its substrates. These conclusions are supported by our findings of overlapping effects of Δ*ctpA* and Δ*lbcA* mutations on the proteome, the occurrence of independent complexes containing LbcA and CtpA or LbcA and substrate *in vivo*, and the direct interaction between purified LbcA and CtpA proteins, and between LbcA and each of the CtpA substrates, *in vitro*. The most abundant LbcA copurification partners were enriched for cell wall enzymes and other cell wall associated proteins. However, some of these LbcA copurification partners are not CtpA substrates, and in contrast to the substrates, those we tested here did not bind to LbcA directly. Their copurification might be explained by indirect association with LbcA, due to their participation in multiprotein complexes containing CtpA substrates. Finally, we have also discovered a novel substrate of the LbcA•CtpA complex, the uncharacterized PA1048 protein.

The LbcA-CtpA system is analogous to the *E. coli* NlpI-Prc system. Both are composed of a TPR-containing outer membrane lipoprotein in complex with a CTP, and both degrade peptidoglycan hydrolases (15–18). Their similarity is further supported by our new finding that LbcA acts as a scaffold, because *in vivo* pull down experiments suggested that NlpI might bind to the Prc protease and its substrates independently as well (16). However, while these two systems are clearly analogous, it is equally clear that they are not orthologous. LbcA is approximately twice the size of NlpI and their primary sequences are not homologous. Furthermore, Prc is a member of the CTP-1 subfamily, whereas CtpA is in the CTP-3 subfamily. In fact, *P. aeruginosa* has a second CTP that is the ortholog of *E. coli* Prc, although little is known about its function. Finally, in other ongoing work we have discovered that the LbcA-CtpA and NlpI-Prc systems work differently at the molecular level, with significant differences in their stoichiometry and structures (H. Hsu, M. Wang, A. Kovach, A. Darwin and H. Li, unpublished data). Therefore, it is intriguing that these two systems have evolved to use quite different components, functioning in different ways, to achieve the common goal of degrading peptidoglycan hydrolases.

We discovered that PA1048 is a substrate of the LbcA•CtpA complex (Fig. 3). In contrast to the four substrates that were already known, PA1048 is not a predicted peptidoglycan hydrolase. However, it does have a link to peptidoglycan because its C-terminal domain, representing approximately 50% of the mature protein, is predicted to adopt an OmpA C-terminal domain fold, which interacts with peptidoglycan non-covalently. This domain occurs in several *P. aeruginosa* proteins, some of which have been demonstrated to interact with peptidoglycan (32–34). Proteins with the OmpA C-terminal domain are widespread in Gram negative bacteria, but their N-terminal domains vary. Some have an N-terminal OM porin domain like OmpA itself, whereas others, including PA1048, are lipoproteins that attach to the OM by their acylated N-termini. The attachment of these proteins to the OM and the cell wall explains how they can affect envelope integrity. For example, *P. aeruginosa* OprF and PA1041 have been linked to cell envelope maintenance (33, 35). A PA1048 null mutant was reported to be sensitive to multiple antibiotics (36). Also, we have preliminary data that a ΔPA1048 mutant has increased sensitivity to SDS/EDTA, which suggests a compromised OM (M. Wang and A. J. Darwin, unpublished data). Global studies suggested that PA1048 gene expression is likely to be under direct control of the AlgR virulence regulator, indicating that PA1048 might be especially important in AlgR-inducing conditions (37, 38). Finally, the four peptidoglycan hydrolase substrates of CtpA are potentially dangerous if their activities are not constrained. Perhaps PA1048 might also be dangerous if its levels are too high at the wrong time. For example, even though PA1048 is not a peptidoglycan hydrolase itself, perhaps it can affect peptidoglycan hydrolase activity indirectly.

In order to conclude that a protein is a protease substrate, it must accumulate in a protease defective mutant *in vivo* and be degraded by the protease *in vitro*. Either one alone is not sufficient because accumulation *in vivo* could be an indirect consequence of absent protease activity, and many proteases can degrade non-physiological substrates *in vitro*. Therefore, even though MltD was degraded by CtpA *in vitro,* it is apparently not a physiological substrate, because MltD did not accumulate in a Δ*ctpA* mutant (Fig. 3). Similarly, only one of ten proteins degraded by *E. coli* Prc *in vitro* was found to accumulate in a Prc-deficient mutant (15). This was not surprising because some of the other nine proteins were cytoplasmic. Perhaps MltD is not degraded because it is physically separated from the LbcA•CtpA complex *in vivo*. For example, if they localize to different regions of the OM. Alternatively, they could be in different membranes altogether. When the sorting signal from *P. aeruginosa* MltD was added to a fluorescent reporter protein, it localized to the inner membrane of *P. aeruginosa* (39). Therefore, *P. aeruginosa* MltD was proposed to be anchored to the inner membrane, which would separate it from the LbcA•CtpA complex. However, these findings have not been corroborated by determining the location of MltD itself, and our attempts to do so have been inconclusive, possibly due to low abundance of MltD and the low avidity of our antiserum. Of course, we cannot rule out the possibility that MltD might be degraded by CtpA *in vivo* under conditions that we are unaware of. However, we have seen no evidence of MltD accumulation in a Δ*ctpA* mutant, including at the different phases of growth.

When LbcA was isolated from *P. aeruginosa* lysates, the most abundant envelope proteins that copurified with it were enriched for cell wall-associated functions (Supplemental Table 6). We followed up on seven of those proteins and found that four of them were not CtpA substrates, the lytic transglycosylases RlpA, MltD and MltB1, and the amidase AmiB. These non-substrates had a striking and unequivocal difference from the CtpA substrates when we tested their ability to bind to LbcA directly *in vitro*. All five known CtpA substrates bound to LbcA, whereas all four of the non-substrates that we tested did not (Fig. 5). A definitive conclusion cannot be made from a negative result, but the uniform ability of all CtpA substrates to bind to LbcA directly, coupled with the failure of all four non-substrates to do so, strongly suggests that these non-substrates co-purified with LbcA due to indirect interactions. This was supported by co-IP experiments, which suggested the occurrence of LbcA-independent complexes *in vivo*, which included CtpA-substrates and non-substrates (Fig. 6). It is also consistent with the known and hypothesized occurrence of multi-enzyme complexes involved in peptidoglycan remodeling during cell growth and division (5, 40). We could not determine which CtpA substrate(s) might bridge interactions between LbcA and the non-substrates. Individual substrate deletion mutations did not prevent RlpA from copurifying with LbcA (Fig. 6 and data not shown) and analysis of multiple substrate deletion mutants has not been possible due to severe growth deficiencies. We also tested if purified RlpA was captured by LbcA *in vitro* if CtpA substrates were also added, but this was not successful. Of course, it is also possible that undiscovered CtpA substrates might be involved.

Does LbcA bind to any proteins directly other than CtpA and its substrates, and does LbcA have any role(s) besides facilitating CtpA-dependent proteolysis? The strikingly similar effects of Δ*ctpA* and Δ*lbcA* mutations on global protein levels suggests that the most impactful role of LbcA is to support proteolysis by CtpA. In addition, global phenotype analysis has shown that *ctpA* and *lbcA* null mutants have very similar phenotypes in several other *Pseudomonas* species (41). However, these analyses do not rule out other roles for LbcA. Furthermore, it is important to note a bias in our study, because the LbcA copurification partners that we focused on here were those that were also cross-linked to an inactive LbcA•CtpA-S302A complex, but not to an active LbcA•CtpA complex, in a previous study (17). This should enrich for protease substrates, and for proteins bound to those substrates, and our follow-up experiments have supported that. Therefore, it remains possible that LbcA has other roles and other direct binding partners beyond CtpA and its substrates. In fact, in the analogous NlpI-Prc system of *E. coli*, evidence suggests that NlpI might have roles beyond facilitating Prc-dependent proteolysis (42). Those authors proposed that NlpI is a general scaffold for peptidoglycan hydrolases, binding to non-Prc substrates and influencing their activity. NlpI and LbcA are not homologous, but both contain TPR repeats that facilitate protein-protein interactions, and so LbcA has the potential to do something similar to NlpI. In fact, NlpI binds to *E. coli* MepS and MepM (although it only promotes degradation of the former), and *P. aeruginosa* LbcA also binds to the MepS homologs PA1198 and PA1199 and to MepM (16, 17, 42). Therefore, NlpI and LbcA have homologous direct binding partners. If LbcA does have direct binding partners that are not CtpA substrates, some obvious candidates would be the cell wall-associated proteins that copurified with LbcA but were not investigated as part of this study (Supplemental Table 6). This will be an interesting topic for future studies. Regardless, even if LbcA only binds to CtpA and its substrates directly, it could still have a role beyond promoting degradation. For example, if there is a population of LbcA in the cell that is not bound to CtpA, but is bound to substrate(s), it could influence their activity or location, and that of any other proteins in complex with those substrates.

In summary, we have shown that LbcA and CtpA have similar impacts on the proteome; that LbcA facilitates CtpA-dependent proteolysis by acting as a scaffold for the protease and its substrates; discovered the first CtpA substrate that is not a peptidoglycan hydrolase; and found evidence of complexes between CtpA substrates and non-substrates *in vivo*. Goals for the future include investigating the significance of complexes containing CtpA substrates and non-substrates; whether or not LbcA can influence the function of these proteins beyond promoting their degradation; and a focused investigation into the possibility that LbcA might have undiscovered direct binding partners beyond CtpA and its proteolytic substrates.

## MATERIALS AND METHODS

### Bacterial strains, plasmids and growth

Strains and plasmids are listed in Table 2. Bacteria were grown in Luria-Bertani (LB) broth, composed of 1% (w/v) tryptone, 0.5% yeast extract, 1% (w/v) NaCl, or on LB agar, at 30°C or 37°C. To select for *P. aeruginosa* exconjugants after mating with *E. coli* donor strains, bacteria were recovered on Vogel-Bonner minimal agar (43).

**TABLE 2.**
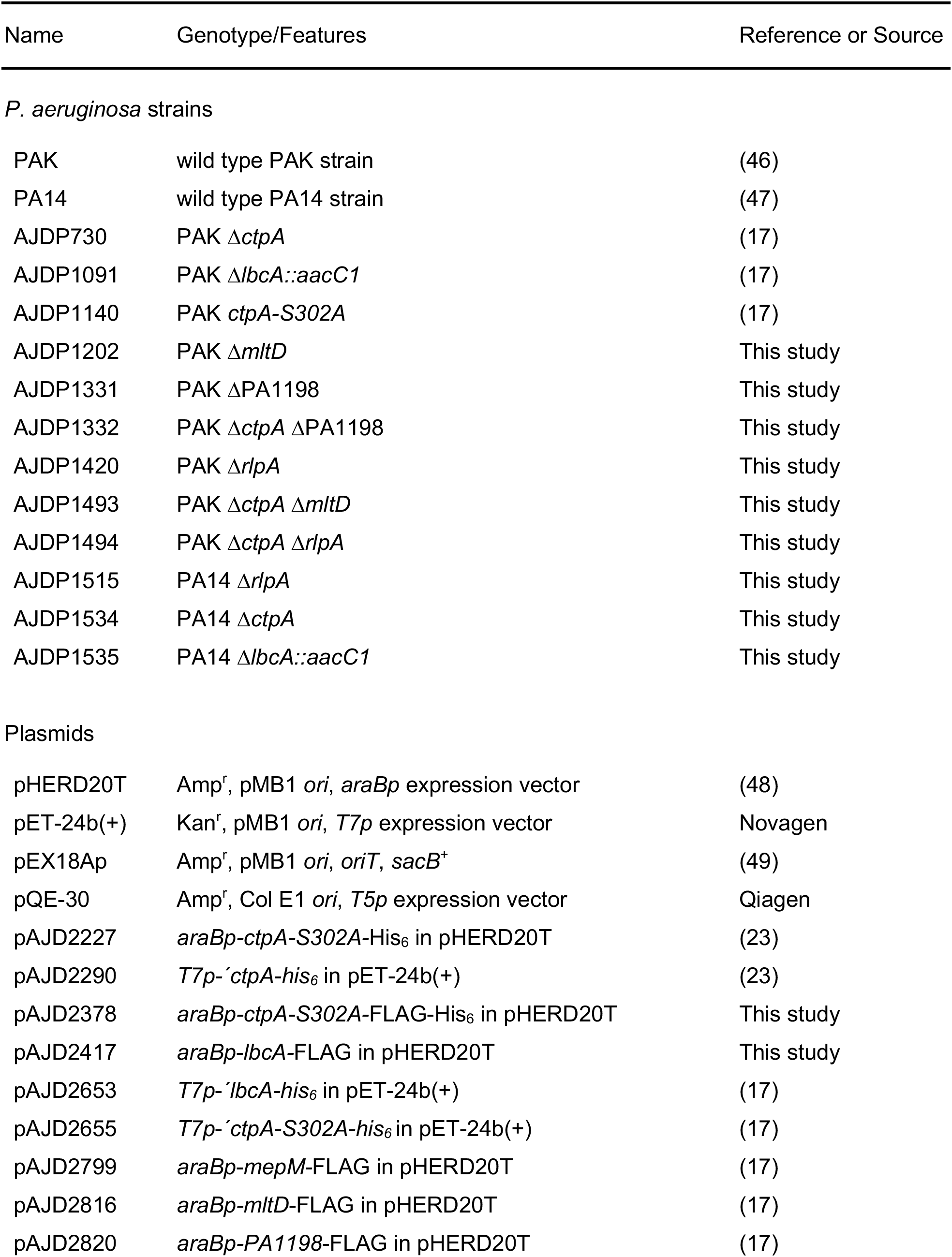
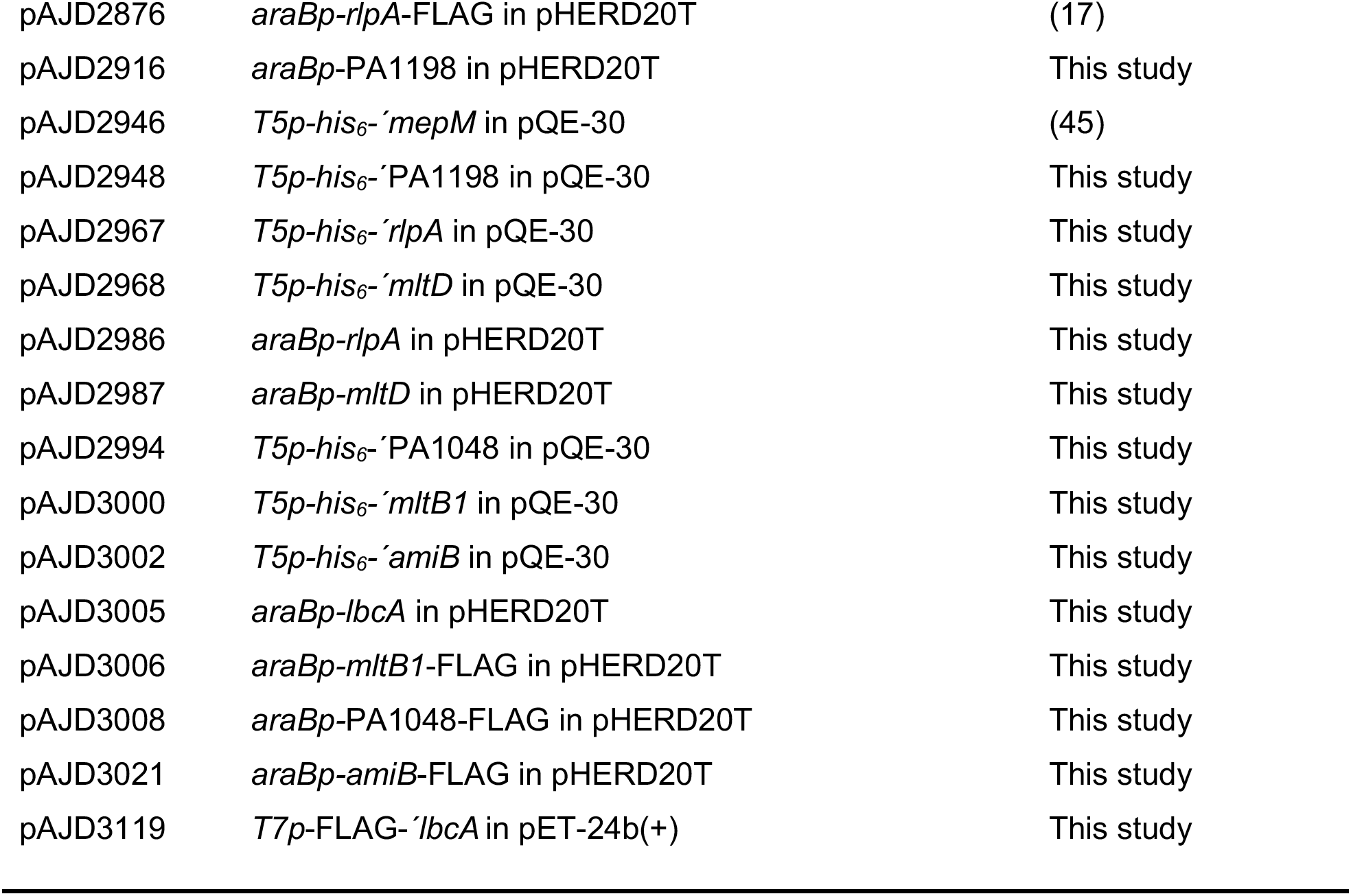
Strains and plasmids

### Plasmid and strain constructions

Construction of strains with Δ*ctpA*, Δ*lbcA::aacC1* and *ctpA-S302A* mutations was described previously (17). To construct strains with ΔPA1198, Δ*mltD* or Δ*rlpA* in frame deletion mutations, two fragments of ~0.55 kb each corresponding to regions flanking the deletion site were amplified by PCR and cloned into pEX18Ap. The plasmids were integrated into the *P. aeruginosa* chromosome after conjugation from *E. coli*, and then sucrose-resistant, carbenicillin-sensitive segregants were isolated on LB agar containing 10% sucrose (44). Deletions were verified by PCR analysis of genomic DNA.

Plasmids encoding C-terminal FLAG- and/or His_6_-tagged proteins were constructed by amplifying the genes from *P. aeruginosa* PAK DNA using a downstream primer that included a region encoding the tag, followed by a stop codon. For plasmids encoding proteins without any added tags, the downstream primer annealed immediately downstream of the stop codon. The amplified fragments were cloned into pHERD20T using restriction sites incorporated by the PCR primers. Plasmids encoding proteins with N-terminal His_6_ tags for protein purification were constructed by amplifying the genes encoding the mature proteins (no N-terminal signal sequence) from *P. aeruginosa* PAK DNA, using a downstream primer that annealed immediately after the stop codon. These fragments were cloned into pQE-30 using restriction sites incorporated by the PCR primers. The plasmid encoding FLAG-′LbcA was constructed by amplifying the region encoding the mature part of LbcA from *P. aeruginosa* PAK DNA, using a forward primer that included a region encoding the FLAG tag and a downstream primer that annealed immediately after the stop codon. This fragment was cloned into pET-24b(+) using NdeI and HindIII restriction sites incorporated by the PCR primers.

### Label-free quantitative proteomics

Bacteria were inoculated into 100 ml of LB broth in a 500 ml flask to an optical density at 600 nm (OD_600_) of 0.05 and grown at 37°C with aeration for 5 h. Cells from the equivalent of 100 ml of culture at OD_600_ = 1 were resuspended in 10 ml of 8 M Urea, 3% (w/v) sodium dodecyl sulfate (SDS), 100 mM Tris-HCl, 1 mM EDTA, pH 8.0, and sonicated. 1 ml was transferred to a microfuge tube and insoluble debris were collected by centrifugation in a microfuge at maximum speed for 20 min. The protein concentration of the clarified lysates, determined with the BioRad DC Protein Assay kit, was approximately 1.2 - 1.5 mg/ml. Triplicate samples were generated for each strain, each from an independent culture. SDS was eliminated with an S-trap column, proteins were digested with trypsin and then identified and quantified by the NYU School of Medicine Proteomics Laboratory using a Thermo Scientific Orbitrap Elite Hybrid Ion Trap-Orbitrap Mass Spectrometer. LFQ values were normalized, filtered for proteins with at least two or more peptides identified, intensity values were Log2 transformed, samples were grouped based on identity (wild type, Δ*ctpA* or Δ*lbcA*), filtered for proteins that were identified in all triplicate samples of at least one strain, missing values were replaced from normal distribution, and then a Student’s t-test was applied followed by a 5% Benjamini-Hochberg based false discovery rate cutoff.

### Anti-FLAG co-immunoprecipitation assay

Bacteria were inoculated into 100 ml of LB broth in a 250 ml flask to an optical density at 600 nm (OD_600_) of 0.05, and grown at 37°C with aeration for 5 h. Equivalent amounts of bacterial cells from all strains were collected by centrifugation, washed in 10 mM Tris-HCl, 10% (w/v) glycerol, pH 7.5 and resuspended in 2 ml non-denaturing lysis buffer (NDLB; 50 mM Tris-HCl, 300 mM NaCl, 5 mM EDTA, 10% (w/v) glycerol, pH 7.5). Roche complete protease inhibitors were added, cells were disrupted by sonication, and then 1% (w/v) Lauryldimethylamine N-oxide (LDAO) was added, followed by incubation with rotation for 1 h at 4°C. Insoluble material was removed by centrifugation at 16,000 x g for 30 min at 4°C. 30 μl of anti-FLAG M2 affinity resin (SigmaAldrich) in NDLB was added to the supernatant and incubated for 2 h at 4°C with rotation. A 1-ml spin column (Pierce; 69725) was used to wash the resin ten times with 500 μl NDLB containing 0.1% (w/v) Triton X-100. Proteins were eluted by addition of 100 μl of 400 μg/ml 3x FLAG peptide in NDLB containing 0.1% (w/v) Triton X-10 and incubation with rotation at 4°C for 30 min. In some cases, proteins present in these samples were identified by liquid chromatography-mass spectrometry (NYU School of Medicine Proteomics Laboratory).

### Polyclonal antisera production and immunoblotting

*E. coli* strain M15 [pREP4] (Qiagen) containing a pQE-30 derivative encoding His_6_-PA1048, His_6_-RlpA or His_6_-MltD was grown in LB broth to mid-log phase at 37°C with aeration. Protein production was induced with 1 mM IPTG for 3 h at 37°C. Proteins were purified under denaturing conditions by nickel-nitrilotriacetic acid (NTA)-agarose affinity chromatography as described by the manufacturer (Qiagen). Polyclonal rabbit antisera were raised by Covance Research Products Inc. Specificity was verified by detection of proteins in whole cell lysates of *P. aeruginosa* strains that either lacked the target protein (deletion mutant), or had it present at endogenous level (chromosomally encoded), or overproduced from a plasmid. Polyclonal antisera for CtpA, LbcA, MepM, PA0943, PA1198 and PA1199 were described previously (17, 23, 30).

Samples separated by SDS-PAGE were transferred to nitrocellulose by semi dry electroblotting. Chemiluminescent detection followed incubation with one of the diluted polyclonal antisera described above, or anti-FLAG M2 monoclonal antibody (Sigma), then goat anti-rabbit IgG (Sigma) or goat anti-mouse IgG (Sigma) horseradish peroxidase conjugates used at the manufacturers recommended dilution.

### Determination of protein abundance *in vivo*

Saturated cultures were diluted into 5 ml of LB broth, containing 150 μg/ml carbenicillin and 0.02% (w/v) arabinose, in 18 mm diameter test tubes so that the OD600 was 0.05. The cultures were grown on a roller drum at 37°C for 5 h. Cells were harvested by centrifugation, resuspended in SDS-PAGE sample buffer at equal concentrations (based on the culture OD600), and analyzed by immunoblot as described above.

### Protein purification

LbcA-His_6_, CtpA-His_6_ and CtpA-S302A-His_6_ were purified exactly as described previously (45). For His_6_-PA1198, His_6_-MepM, His_6_-MltD, His_6_-RlpA and His_6_-AmiB, *E. coli* strain M15 [pREP4] containing a pQE-30 plasmid derivative was grown in 500 ml LB broth at 37°C with aeration to an OD_600_ of 0.6 - 1.0. Protein production was induced by adding 1 mM IPTG and incubation at 37°C was continued for 3 h, after which cells were harvested by centrifugation. Proteins were purified under native conditions by NTA-agarose affinity chromatography in buffer containing 50 mM NaH_2_PO_4_ and 300 mM NaCl, as recommended by the manufacturer (Qiagen). Proteins were eluted in 1 ml fractions using 50 mM NaH_2_PO_4_, 300 mM NaCl, pH 8 buffer containing 50, 100, 150, 200, or 250 mM imidazole.

His_6_-PA1048 and His_6_-MltB1 encoded by pQE-30 plasmid derivatives were purified differently because they were insoluble when overproduced in *E. coli*. Proteins were denatured and solubilized by resuspending the harvested cells in 6M Guanidine hydrochloride, 100 mM NaH_2_PO_4_, 10 mM Tris-HCl, 5 mM imidazole, 1% (w/v) Triton X-100, pH 7.5, and stirring at room temperature for 1 h. The suspension was sonicated and insoluble material was removed by centrifugation at 10,000 × *g* for 30 min. Ni-NTA agarose was added to the supernatant and it was stirred for 1 h at room temperature. The Ni-NTA agarose resin was collected in a drip column and washed with 30 ml of 100 mM NaH_2_PO_4_, 10 mM Tris-HCl, 1% (w/v) Triton X-100, 20 mM imidazole, 8M Urea, pH 7.5. Proteins were renatured by washing the resin with 50 ml of 50 mM NaH_2_PO_4_, 300 mM NaCl, 20 mM imidazole, pH 8. Proteins were eluted in 1.5 ml fractions using 50 mM NaH2PO4, 300 mM NaCl, pH 8 buffer containing 50, 100, 150, 200, or 250 mM imidazole. His_6_-RlpA was also repurified this way to demonstrate that the denaturation/renaturation process would not render a non-CtpA substrate susceptible to cleavage.

### *In vitro* proteolysis assay

These were done similarly to those described previously (17, 45). N-terminal His_6_-tagged test proteins were mixed with CtpA-His_6_ or CtpA-S302A-His_6_, together with LbcA-His_6_, all purified as described above, and incubated at 37°C for 3 h. Reactions were terminated by adding SDS-PAGE sample buffer and incubating at 90°C for 10 min. The samples were separated by SDS-PAGE and stained with ProtoBlue Safe (National Diagnostics).

### LbcA direct binding assay

*E. coli* strain ER2566 (New England Biolabs) containing a pET-24b(+) derivative encoding FLAG-LbcA was grown in 250 ml LB broth at 37°C with aeration to an OD_600_ of approximately 0.6. Production of FLAG-LbcA in the *E. coli* cytoplasm was induced by adding 1 mM IPTG and incubation at 37°C was continued for 3 h, after which cells were harvested by centrifugation. Cells were washed in 10 mM Tris-HCl, 10% (w/v) glycerol, pH 7.5 and resuspended in 5 ml NDLB. Roche complete protease inhibitors were added, cells were disrupted by sonication, and then 1% (w/v) LDAO was added, followed by incubation with rotation for 1 h at 4°C. Insoluble material was removed by centrifugation at 16,000 × g for 30 min at 4°C. A final concentration of 5% (w/v) glycerol was added to the supernatant, which was stored in aliquots at −70°C (this was the FLAG-LbcA soluble lysate). A negative control lysate was generated the same way using *E. coli* ER2566 containing the empty vector plasmid pET-24b(+).

For direct binding assays, purified baits were prepared by adding 10 μl of EZview™ Red anti-FLAG M2 affinity resin (SigmaAldrich) in NDLB to 50 μl of either the FLAG-LbcA soluble lysate, or the negative control soluble lysate, and the total volume was adjusted to 0.5 ml with NDLB. This mixture was incubated with rotation at 4°C for 2 h, the resin was collected by centrifugation, and then washed five times with 0.5 ml high salt NDLB (NDLB containing 0.5 M NaCl) containing 0.1% (w/v) Triton X-100. The purified anti-FLAG M2 agarose immunocomplex was blocked by resuspending in 0.5 ml high salt NDLB containing 5% (w/v) bovine serum albumin, incubating with rotation at 4°C overnight, and then washing the resin five times with 0.5 ml high salt high salt NDLB containing 0.1% (w/v) Triton X-100. To test for direct binding, the resin was resuspended in 0.5 ml high salt high salt NDLB containing approximately 1.5 μg of a His_6_-tagged prey protein, which had been purified as described above. Binding was allowed to occur by incubating with rotation at 4°C for 1 h and the resin was collected by centrifugation, washed five times with 0.5 ml high salt high salt NDLB containing 0.1% (w/v) Triton X-100, resuspended in 100 μl SDS-PAGE sample buffer and boiled for 10 min to elute proteins. 10 μl of these samples was loaded into each gel lane for SDS-PAGE and immunoblot analysis.

## ACKNOWLEDGEMENTS

Research was supported by the National Institute of Allergy and Infectious Diseases (NIAID) of the National Institutes of Health, under Award Number R01AI136901. The content is solely the responsibility of the authors and does not necessarily represent the official views of the National Institutes of Health. Mass spectrometric protein identification done by NYU Grossman School of Medicine’s proteomics laboratory was partly supported by NYU Grossman School of Medicine.

We thank Alexis Sommerfield for providing critical comments on a draft version of the manuscript.

